# Unveiling the Catalytic Mechanism of GTP Hydrolysis in Microtubules

**DOI:** 10.1101/2023.05.01.538927

**Authors:** Daniel Beckett, Gregory A. Voth

## Abstract

Microtubules (MTs) are large cytoskeletal polymers, composed of αβ-tubulin heterodimers, capable of stochastically converting from polymerizing to depolymerizing states and vice-versa. Depolymerization is coupled with hydrolysis of GTP within β-tubulin. Hydrolysis is favored in the MT lattice compared to free heterodimer with an experimentally observed rate increase of 500 to 700 fold, corresponding to an energetic barrier lowering of 3.8 to 4.0 kcal/mol. Mutagenesis studies have implicated α-tubulin residues, α:E254 and α:D251, as catalytic residues completing the β-tubulin active site of the lower heterodimer in the MT lattice. The mechanism for GTP hydrolysis in the free heterodimer, however, is not understood. Additionally, there has been debate concerning whether the GTP-state lattice is expanded or compacted relative to the GDP-state and whether a “compacted” GDP-state lattice is required for hydrolysis. In this work, extensive QM/MM simulations with transition-tempered metadynamics free energy sampling of compacted and expanded inter-dimer complexes, as well as free heterodimer, have been carried out to provide clear insight into the GTP hydrolysis mechanism. α:E254 was found to be the catalytic residue in a compacted lattice, while in the expanded lattice disruption of a key salt bridge interaction renders α:E254 less effective. The simulations reveal a barrier decrease of 3.8 ± 0.5 kcal/mol for the compacted lattice compared to free heterodimer, in good agreement with experimental kinetic measurements. Additionally, the expanded lattice barrier was found to be 6.3 ± 0.5 kcal/mol higher than compacted, demonstrating that GTP hydrolysis is variable with lattice state and slower at the MT tip.

**Significance Statement:** Microtubules (MTs) are large and dynamic components of the eukaryotic cytoskeleton with the ability to stochastically convert from a polymerizing to a depolymerizing state and vice-versa. Depolymerization is coupled to the hydrolysis of guanosine-5’-triphosphate (GTP), which is orders of magnitude faster in the MT lattice than in free tubulin heterodimers. Our results computationally ascertain the catalytic residue contacts in the MT lattice that accelerate GTP hydrolysis compared to the free heterodimer as well as confirm that a compacted MT lattice is necessary for hydrolysis while a more expanded lattice is unable to form the necessary contacts and thereby hydrolyze GTP.

## Introduction

Microtubules (MTs) are ubiquitous components of the eukaryotic cytoskeleton that play key roles in a plethora of cellular processes, including serving as tracks for motor proteins and creation of the dynamic mitotic spindle during mitosis.^1-3^ A specific property of MTs that allows them to fulfill such diverse roles is dynamic instability (DI), wherein growing MTs stochastically switch from a polymerizing state to a depolymerizing state (MT catastrophe) and vice-versa (rescue).^4-6^ The delicate balance of the DI process makes the MT a potent drug target to treat multiple cancers through the manipulation of MT structure and catalytic activity, in order to either discourage or encourage depolymerization.^7-10^

MTs are assembled from tubulin heterodimers, each composed of an α- and β-tubulin monomer, which stack into protofilaments (PFs) and laterally associate into tubules. While both monomeric tubulin subunits bind a molecule of guanosine triphosphate (GTP), in the free heterodimer β-tubulin sits atop the GTP binding site of α-tubulin while in the microtubule lattice the α-tubulin of one heterodimer sits atop the GTP binding site of another heterodimer’s β-tubulin (Fig. 1A–C). GTP bound by α-tubulin is not hydrolyzed or exchanged in solution with other GTP molecules, whereas GTP bound by β-tubulin can hydrolyze to guanosine diphosphate (GDP) and subsequently GDP can be exchanged at the microtubule tip, or in the free heterodimer, for GTP. These observations have led to the α-tubulin GTP binding site to be called the non-exchangeable site (N-site) while β-tubulin possesses the exchangeable site (E-site).^11-14^ While it is known that the hydrolysis of GTP at the E-site is coupled to microtubule depolymerization and catastrophe, the exact mechanism behind this is not fully understood and insights into the actual hydrolysis mechanism would be invaluable.^15,16^

**Figure 1.**
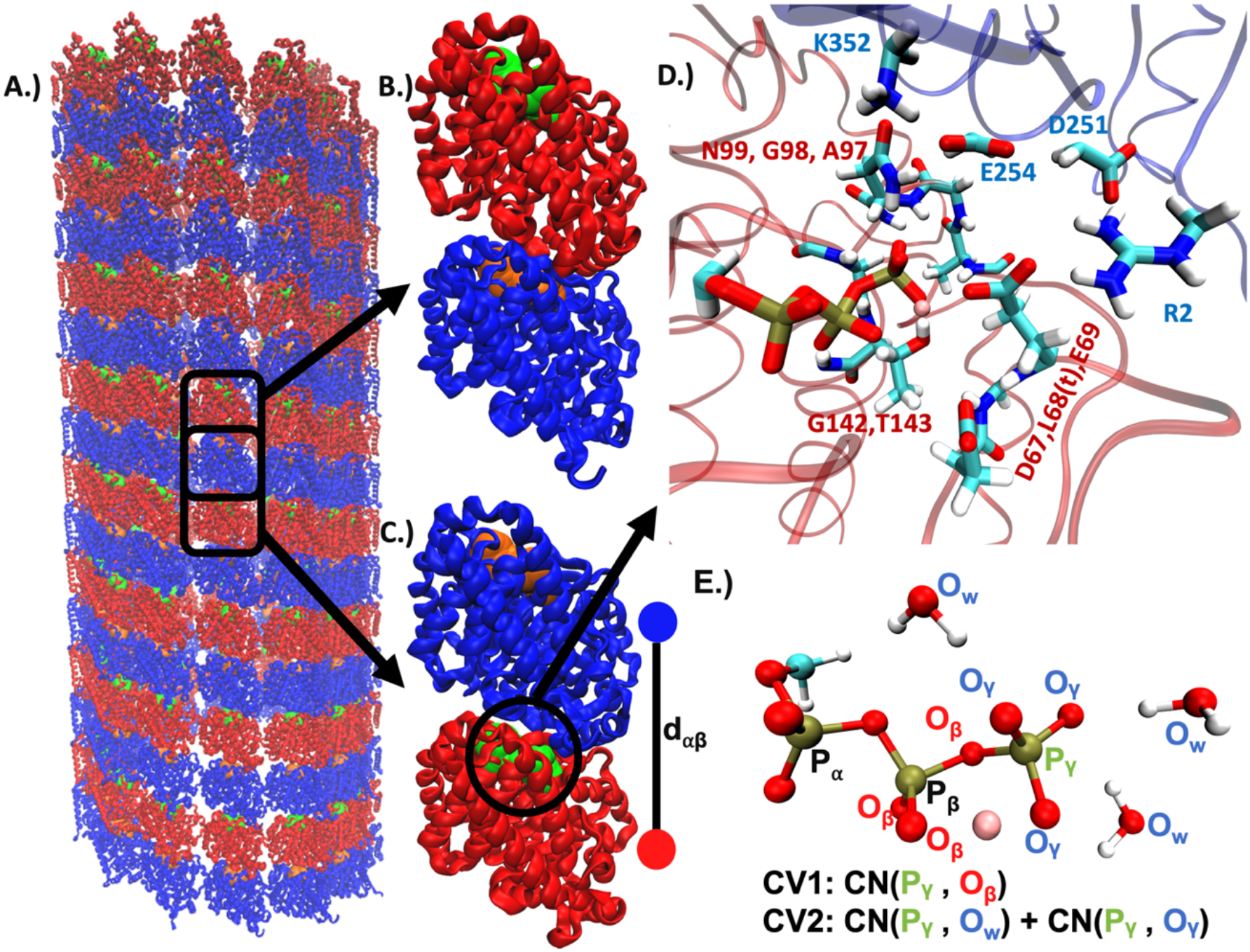
Schematics of systems studied: A.) cartoon of a full microtubule, plus-end on top, with β-tubulin in red, α-tubulin in blue, GTP bound by β-tubulin in green, GTP bound by α-tubulin in orange, B.) example of a tubulin αβ-heterodimer, C.) example of an inter-dimer complex formed by the α-tubulin of one heterodimer sitting above the β-tubulin of another, with d_αβ_ defined as the distance between the centers of mass of the monomers, D.) the QM region used in this study for inter-dimer complexes (more description in Methods), with magnesium in pink, E.) labeled atoms for reaction coordinate definitions as seen in Figure 2: x-axis is the coordination number of P_γ_ to O_β_, y-axis is the coordination number of P_γ_ to O_γ_ plus the coordination number of P_γ_ to O_w_.

GTP hydrolysis has been known to be faster in the MT lattice than the free heterodimer, with recent experiments anticipating a rate difference of 500 to 700 fold, corresponding to an energetic barrier difference of 3.9 ± 0.1 kcal/mol.^17,18^ Cryo-EM measurements and mutagenesis studies on have led to the hypothesis that specific α-tubulin residues (α:E254 and α:D251) complete the β-tubulin GTPase site and catalyze hydrolysis.^18-22^ Additionally, when a non-hydrolyzable GTP-mimic, GMPCPP, is used in place of GTP the MT lattice seems to be expanded relative to the GDP lattice, which supports the hypothesis that lattice compaction induced by hydrolysis incurs strain on the MT lattice, which in turn is released by catastrophe.^23,24^ Recent work using inorganic phosphate mimics has implied that expansion is an artifact of GMPCPP in the lattice which would also imply that lattice compaction is necessary for hydrolysis rather than a result of it.^25^ Finally, recent cryo-EM analysis of mutated (α: E254A/N) MT has revealed that the expanded GMPCPP state may more closely resemble tubulin at the MT tip.^26^

The present study aims to deconvolute the discussion of microtubule structure by investigating GTP hydrolysis both in an expanded and compacted MT lattice, as well as in the free heterodimer. Through the use of transition-tempered metadynamics (TTMetaD) enhanced free energy sampling combined with quantum mechanices/molecular mechanics (QM/MM) simulation, we have mapped the free energy surface of GTP hydrolysis and affirmed the importance of the α:E254 residue in the catalytic activity of the compacted lattice. We have determined that the activation barrier of hydrolysis in the free heterodimer is 3.8 ± 0.5 kcal/mol higher than in the compacted lattice, in good agreement with the experimental value of 3.9 ± 0.1 kcal/mol. Furthermore, hydrolysis in the expanded lattice was found to have a higher barrier than even the free heterodimer, potentially due to disruption of a critical inter-subunit salt bridge interaction making α:E254 unreachable. These explicit simulation results confirm the catalytic role of the α:E254 residue while also demonstrating that lattice compaction is essential for facile GTP hydrolysis.

## Results and Discussion

### A Compacted Inter-dimer MT Lattice Hydrolyzes GTP More Readily Than an Expanded Lattice or Free Heterodimer

GTP hydrolysis was explored in 3 systems as illustrated in Figure 1A–C: a free tubulin heterodimer, an inter-dimer complex restrained at a “compacted” GDP-like inter-dimer distance (d_αβ_ = 40.5 Å), and an “expanded” inter-dimer complex held at an inter-dimer distance close to that observed in the tip of an MT (d_αβ_ = 42.0 Å). In all cases, 10 replicas were employed for QM/MM TTMetaD along 2 collective variables (CVs) to define hydrolysis. The final two-dimensional free energy surfaces for GTP hydrolysis in each system can be seen in Figure 2. The QM region and employed CVs can be seen in Figure 1D and 1E, respectively, with additional details in presented in the Materials and Methods.

**Figure 2.**
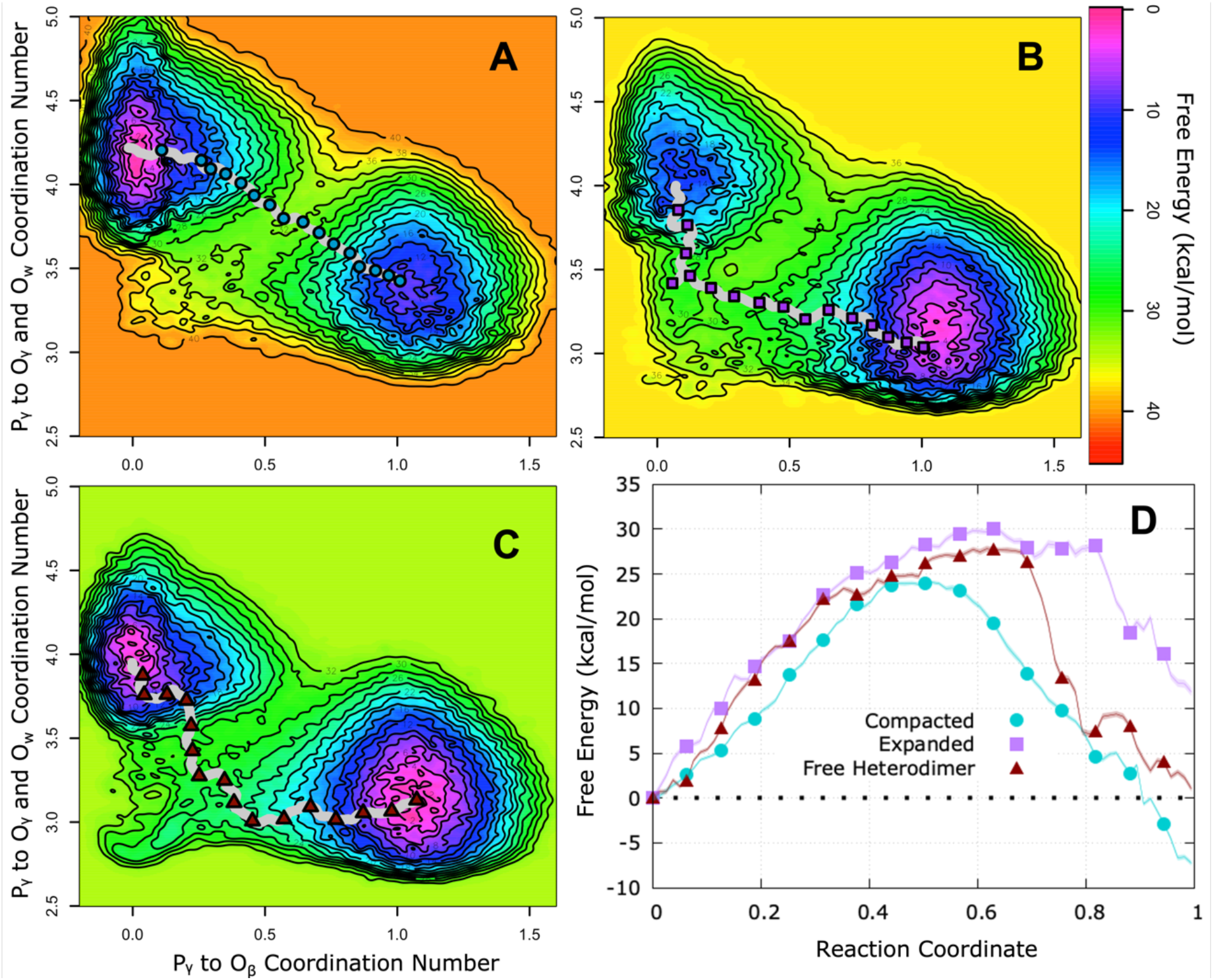
Final converged free energy surfaces for GTP hydrolysis in each system with lowest energy pathway shown as a grey line with symbols overtop: A.) compacted inter-dimer complex, B.) expanded inter-dimer complex, C.) free heterodimer, and D.) the free energy along the lowest energy pathway in each system with error shown as the transparent shading around each line. Topological lines in A – C drawn every 2 kcal/mol. Each pathway is represented by 160 points in total, every 10^th^ point represented as a visible point on the plot for comparison. The x-axis in 2D scales progress along the reaction path from 0 (reactant well) to 1 (product well).

In Figure 2, the *x*-axis represents the coordination number of the leaving dissociating phosphate phosphorus atom to the GDP end phosphate oxygens and varies from ∼ 1 in the reactant well to ∼ 0 in the product well. The *y*-axis shows the coordination number of the phosphorus atom to its own phosphate oxygens as well as any water oxygens and varies from ∼ 3 in the reactant well to ∼ 4 in the product well. A “fully associative” path from reactant to product would be a near vertical line from the reactant well followed by a near horizontal line to the products, whereas a “fully dissociative” path would be a nearly horizontal line from the reactant well followed by a nearly vertical line to products. A diagonal line from reactant to product would be a more concerted path wherein dissociation from GDP is coupled to association of water, similar to that observed in ATP hydrolysis in actin and likely accelerated by a catalytic residue.^27,28^ In the case of the compacted inter-dimer complex a concerted path was found to be the lowest free energy pathway connecting reactants to products, while in the expanded inter-dimer complex and the free heterodimer, a more dissociative pathway was preferred. In both the free heterodimer and the expanded inter-dimer complex, a concerted mechanism was also found but higher in free energy than the dissociative route. Discussion of other possible pathways can be found in the Supplementary Information.

The energetics along the minimum free energy paths (MFEPs) are shown in Figure 2D with progress along the pathway interpolated from 0 (reactant well) to 1 (product well). It is clear from Figure 2D that the lowest energy transition state for any system is along the concerted pathway in the compacted inter-dimer complex, at 24.0 ± 0.2 kcal/mol. As was the case in actin, the barriers are somewhat higher than that expected from experimental rates given the known error in the electronic density functional (DFT) utilized in the QM/MM, but we are primarily interested in the differences between the pathways as that error then cancels out.^27,28^ The barrier for the free heterodimer MFEP is 3.8 ± 0.5 kcal/mol higher than compacted while the barrier for the expanded inter-dimer complex is 6.3 ± 0.5 kcal/mol higher than compacted. The rate difference between the compacted and free heterodimer concerted pathways corresponds well to recent experimental measurements of GTP hydrolysis in a MT (0.16 s^−1^) versus free tubulin (0.0003 s^−1^), which translates to an energetic difference of 3.9 kcal/mol.^17^ Another recent study comparing wildtype MTs to E254D mutants determined a rate of 0.2 s^−1^ for GTP hydrolysis in an MT, corresponding to an energy difference with the free tubulin measurement of 4.00 kcal/mol.^18^ Comparing these experimental values gives us a kinetic barrer of 3.9 ± 0.1 kcal/mol, which is to within error excellent agreement with our compacted vs. free heterodimer concerted pathway barrier difference of 3.8 ± 0.5 kcal/mol. This agreement, as well as the high barrier for hydrolysis in the expanded inter-dimer complex, confirms that lattice compaction is necessary for GTP hydrolysis in an MT.

### αE254 is the Catalytic Residue in the Compacted Lattice, but Unreachable in Expanded

Full movies displaying each type of reaction (concerted and dissociative) for each system can be seen in the Supplementary Information along with the replica ID of the TTMetaD trajectory from which it was taken. Complete 2 CV trajectories overlayed over the free energy surfaces shown in Figure 2 for each replica in each system can also be found in the Supplementary Information.

Figure 3 shows representative snapshots for the minimum free energy pathway in each system, taken directly from the appropriate trajectory. The first panel of each row shows the initial proton transfer from the lytic water. In the compacted inter-dimer complex (Figure 3A), α:E254 serves as the catalytic residue and initiates the deprotonation. This is in line with mutagenesis experiments in both human tubulin and yeast that found an α:E254A mutation to prohibit GTP hydrolysis.^18,21^ In every observed crossing in the compacted inter-dimer complex, α:E254 always fills this role and always does so directly without a water wire being necessary. After the initial deprotonation, the proton is deposited onto an O_γ_ (3A, panel 2) to form the final H_2_PO_4_ product (3A, panel 3).

**Figure 3.**
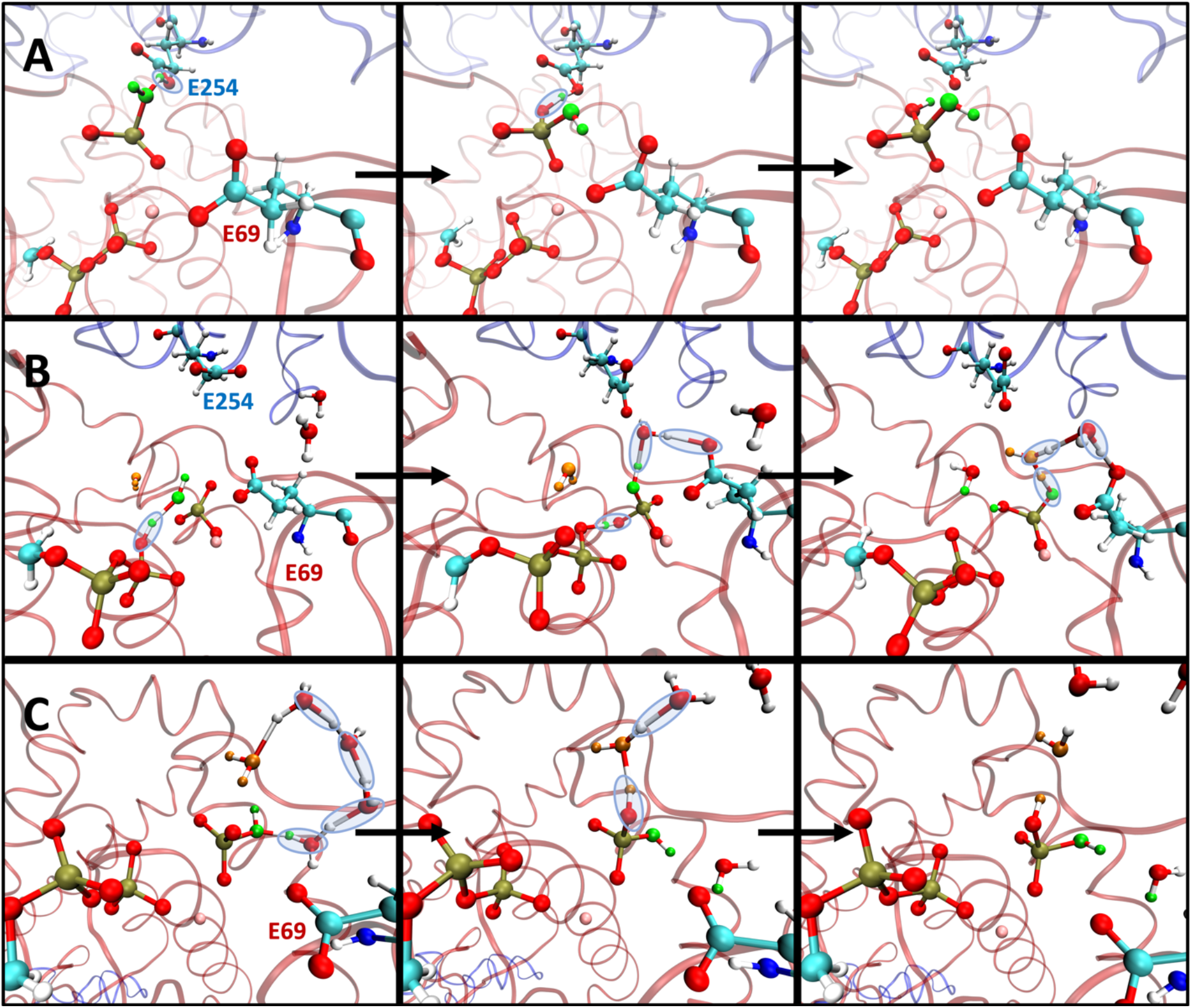
Snapshots of proton rearrangements from metadynamics trajectories for lowest energy reaction pathways in A.) compacted inter-dimer complex (taken from replica ID “W0”) B.) expanded inter-dimer complex (W0), and C.) free heterodimer (W0). Lytic water associating with leaving phosphate is colored green, additional water depositing proton onto the phosphate is colored orange when present (B and C), magnesium shown in pink. Bonds drawn with a 1.8 Å cutoff, only critical residues shown atomically (labeled on the first slide), red ribbons represent β-tubulin while blue represent α-tubulin. The first column shows initial proton transfer, the second column is proton transfer to form H_2_PO_4_, and the third column shows final product after formation (A and C) or final proton transfer (B). Blue translucent ovals highlight proton transfers: circled atoms generally remain bonded after the given frame, though in C this is difficult to define due to the long chain of proton transfers in panels 1 and 2, which occur in quick succession. Movies pulled directly from the appropriate trajectory for each mechanism shown here can be found in the Supplementary Information.

In the free heterodimer (Figure 3C), where the α-tubulin residues are unavailable, PO_3_^−^ dissociates far enough from the GDP to form a water wire with itself that deprotonates the associated H_2_O (3C, panel 1) and protonates another O_γ_ (panel 2) to form H_2_PO_4_. This mechanism results from the trajectory that most closely matches the MFEP shown in Figure 2C; however other higher energy dissociative crossings, similar to the expanded inter-dimer complex, were observed. In the expanded lattice (Figure 3B), presence of the α-tubulin prevents the water wire mechanism seen in the free heterodimer and instead a β phosphate oxygen serves to initiate deprotonation as the lytic water inserts between the β and γ phosphates. In the subsequent panel, β:E69 coordinates the hydroxyl group via a second water and abstracts the second proton while the original proton on the O_β_ is deposited onto a second O_γ_. In the third panel, a final proton transfer is shown where the proton from the orange water is deposited onto the lytic water oxygen through another water wire interaction with β:E69. A second, more direct, mechanism was seen in the expanded inter-dimer complex that would follow the same MFEP, wherein the lytic water was simultaneously coordinated by O_β_ and β:E69 before deprotonation by O_β_. Additionally, concerted pathways involving deprotonation of the lytic water by β:E69 were seen in both the free heterodimer and expanded inter-dimer complex. These pathways are slightly higher in energy (< 1 kcal/mol) than the dissociative pathways and worth consideration as possible mechanisms in the less favorable systems. All alternate mechanisms and free energy paths are documented in the Supplementary Information.

Interestingly, there were no observed TTMetaD barrier crossings in the expanded inter-dimer complex that resulted from deprotonation by α:E254. Occasionally α:E254 is close enough to the lytic water to loosely coordinate a hydrogen (as can be seen in Figure 3B, panel 2), but it does not directly participate in hydrolysis. To understand why α:E254 is unable to serve as the catalytic residue in the expanded inter-dimer complex, Figure 4 compares geometric parameters within the E-site between the compacted and expanded conformations. Most immediately observable is the change in the angle formed by the phosphorus atoms (P_α_–P_β_–P_γ_) in the GTP chain, illustrated by the green line in Figure 4A. This difference in phosphate chain angle is quantified in Figure 4G, comparing the normalized probability distribution over the initial 50 ps metadynamics seed trajectory (a diagram of these trajectories, and all trajectories, are available in the Supplemental Information), the compacted phosphate chain is able to sample a range of more obtuse angles putting it in line with the α:E254 carboxylate group.

**Figure 4.**
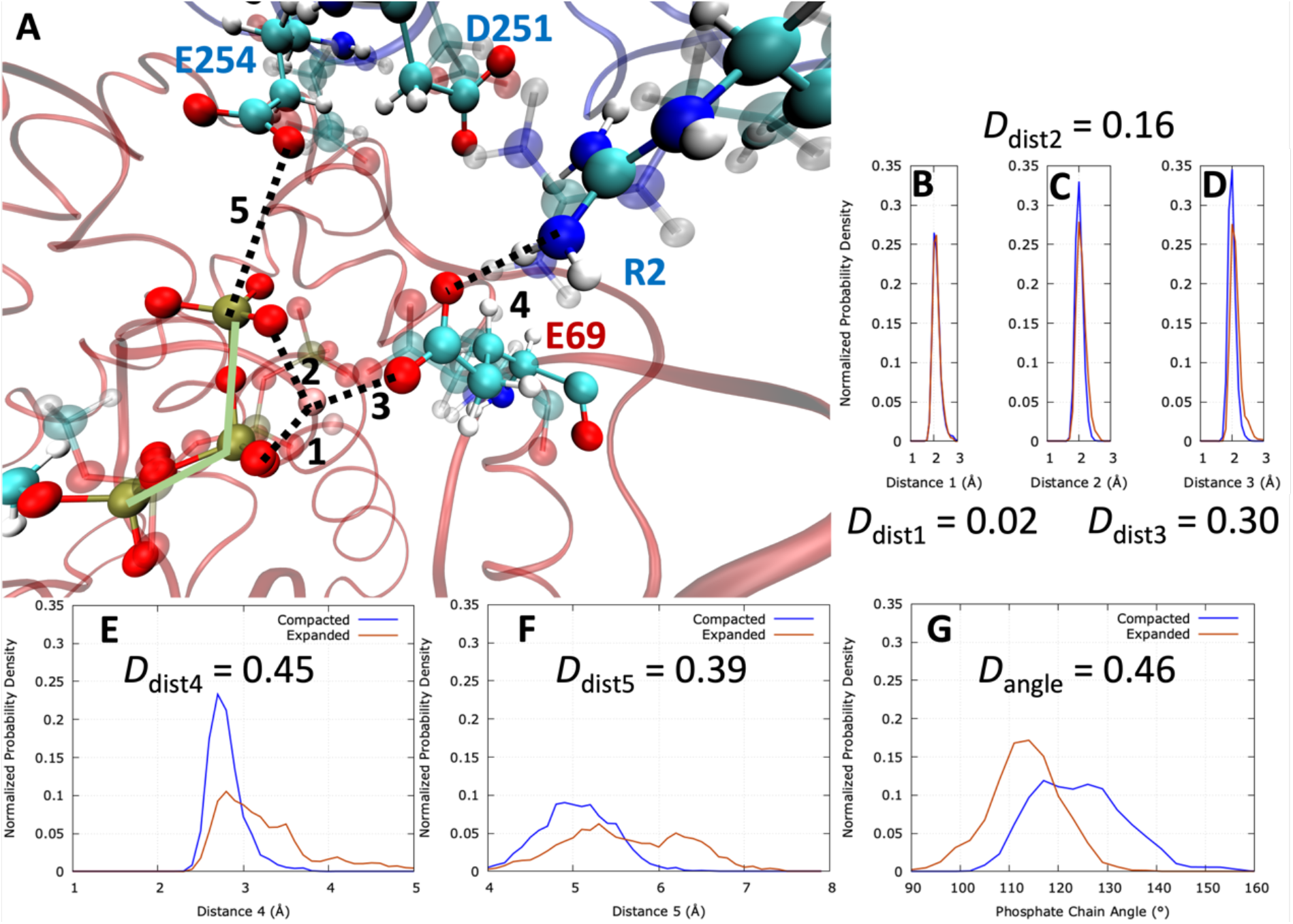
Comparisons of expanded vs. compacted inter-dimer complex geometric parameters within the E-site. A.) CPK rendering of compacted (solid) and expanded (transparent) residues with the compacted protein environment shown as transparent ribbons, taken from the first frame of metadynamics simulation. B–F: Normalized probability distributions of the distances labeled in 5A, computed over the 50 ps reaction well “seed trajectories”. G: Normalized probability distribution of the phosphate chain angle (P_α_-P_β_-P_γ_) over the seed trajectories, illustrated as a green line in 5A. Kolmogorov–Smirnov statistic (D) between the expanded and compacted distributions shown for each plot. Labeled distances: **1** nearest O_β_ to Mg, **2** nearest O_γ_ to Mg, **3** nearest β:E69 carboxylate O to Mg, **4** nearest β:E69 carboxylate O to α:R2 guanidinium N, **5** P_γ_ to nearest α:E254 carboxylate O.

To quantify the difference between the normalized probability distributions of the compacted complex and the expanded complex, we employ the Kolmogorov–Smirnov statistic, *D*, a measure of distance between two probability distributions which is defined as the maximum absolute difference in the cumulative distribution functions.^29,30^ A value of 0 would correspond to identical distributions while a value of 1 (when applied to normalized distributions) would correspond to distributions with no overlap whatsoever. A total of 5 critical contacts are labeled in Figure 5A: the two involving the negatively charged β and γ phosphate oxygen atoms and the Mg^2+^ ion (5B and 5C) have the lowest KS statistics. In both complexes, the β:E69 residue coordinates Mg^2+^ and the expanded complex experiences slightly higher lengths of this interaction (5D); however, it is the inter-dimer interaction of α:R2 and β:E69 (between the positively charged guanidinium N and the negatively charged carboxylic acid O) that displays the highest KS statistic (5E). While this distance in the compacted complex peaks strongly at 2.7 Å, in the expanded complex the α:R2 to β:E69 distance meanders and cannot form steadily. In the compacted complex, the low distance between the R and E heavy atoms qualifies as a complex salt bridge interaction but the interaction is degraded in the expanded complex.^31,32^ Degradation of the α:R2–β:E69 interaction is more pronounced than any other in the immediate E-site binding pocket and likely corresponds to the dip in the phosphate chain angle (5G) and the increase in the critical α:E254–P_γ_ distance (5F) that allows α:E254 to function as a catalytic residue. Mutagenesis of α:R2 to either a complete deactivation (α:R2A) or damping the interaction (α:R2K) would be an interesting comparison in the future, to determine if this has an effect on the rate of hydrolysis in a growing MT.

## Conclusions

GTP hydrolysis in a microtubule is coupled to the onset of depolymerization. The regulation of this reaction is essential to the operation of microtubule associating proteins and drugs targeting their function. Structurally, the hydrolysis of GTP to GDP has been suggested to result in lattice compaction with the nonhydrolyzable GMPCPP mimic and the microtubule tip structure displaying an expanded lattice. Furthermore, the E254 and E251 residues of the α tubulin belonging to the heterodimer above a given β-tubulin-bound GTP have been implicated by mutagenesis experiments as critical to hydrolysis.^18,21^ Extensive QM/MM TTMetaD simulations on a compacted inter-dimer tubulin complex, an expanded inter-dimer tubulin complex, and a free tubulin heterodimer have been reported in this work in order to untangle the effects of compaction and confirm the identity of certain catalytic residues.

Ten TTMetaD QM/MM replicas were run for each system, for a total of ∼10 M cpu*hours of simulation time. In the compacted inter-dimer complex, a concerted pathway is strongly favored and α:E254 directly participates in hydrolysis. In the expanded inter-dimer complex, the triphosphate chain is unable to coordinate a water close enough to the α:E254 carboxylate for it to aid in hydrolysis. This effect may be a result of an inter-dimer salt bridge between α:R2 and β:E69 that is persistent in the compacted lattice but is not fully established in the expanded.

In both the free heterodimer and the expanded inter-dimer complex, a more dissociative pathway was found where the lytic water was deprotonated by the β phosphate oxygens either directly or through a water wire. In both the expanded lattice and the free heterodimer, the dissociative route was somewhat more favorable than a concerted route using the inconveniently located β:E69 residue. The lowest barrier difference between free heterodimer and the compacted inter-dimer complex is 3.8 ± 0.5 kcal/mol, corresponding very well to the experimental value of 3.9 ± 0.1 kcal/mol. Lastly, the expanded inter-dimer complex barrier is 6.3 ± 0.5 kcal/mol higher than the compacted case, confirming that a compacted lattice is essential for facile GTP hydrolysis in a MT and that α:E254 serves as the catalytic factor in the compacted lattice. Furthermore, given the recent cryo-EM measurements of an expanded MT tip,^33^ these results imply GTP hydrolysis proceeds very slowly in the tip compared to within the interior region of the MT.

## Materials and Methods

### System Setup and Equilibration

The initial structures for free heterodimer and the inter-dimer complex were pulled from the 150 ns point on a previous full MT MD simulation of an 8-dimer long 13-PF MT.^34^ A single β-tubulin was chosen from the “back” PF of the MT (opposite to the seam) and the middle layer. For the inter-dimer complex, the α-tubulin about this β-tubulin was extracted while for the free heterodimer the α-tubulin below was extracted. This was done to ensure the same initial β-tubulin GTP binding site configuration for each system. Each system was initially equilibrated in the NVT ensemble for 100 ns in a 121.5 Å side-length cubic periodic box (124.4 Å for the free heterodimer) with 100 mM NaCl at 310 K using the CHARMM36m force field and GROMACS 2019.4.^35,36^ Time step for equilibration was 2 fs and smooth particle mesh Ewald electrostatics were used for the MM throughout both this step and the QM/MM to follow.^37^

The free heterodimer was able to relax without restraints while the inter-dimer complex was restrained at either the compacted or expanded C_α_ and C_β_ center-of-mass (CoM) distance between the α- and β-tubulin monomers with a harmonic force constant of 59.75 kcal/molÅ^2^. The force constant was chosen that reproduced fluctuations in the inter-dimer distance most closely to those observed in the MT tip simulations.^34^ The compacted αβ CoM distance, 40.5 Å was determined from the atomic model cryo-EM structure of an undecorated GDP lattice patch with accession code 6DPV.^24^ The expanded αβ CoM distance, 42.0 Å was derived from the average over all inter-dimer distances in the “back” lattice patch of the full MT tip simulations (PFs 5 through 9 when counting from the seam and dimers 3 through 6) over the last 100 ns of the 200 ns total trajectories.^34^

### QM/MM Setup

Enhanced free energy sampling of the GTP hydrolysis in each system were simulated using the QM/MM hybrid technique as implemented in CP2K version 6.1 with a 0.5 fs timestep.^38-41^ The MM environment was at the same level as described in the previous section. The QM high level used was the PBE density functional, with D3 dispersion corrections and zero damping, using the TZV2P basis set and GTH pseudopotential for the magnesium ion core electrons.^42-47^ All simulations were run in the NVT ensemble with a 121.5 Å side-length cubic periodic box (124.4 Å for the free heterodimer), 100 mM NaCl, and at 310 K using a Nosé-Hoover thermostat.

The quantum region for the inter-dimer complexes included the GTP truncated as a methyl triphosphate, the associated magnesium atom, waters within 6 Å of the γ-phosphate (7 Å for the free heterodimer), and 8 β-tubulin and 4 α-tubulin amino acids near the phosphate tail. β-tubulin amino acids included in the QM region were Asp67, Leu68 (sidechain truncated), Glu69, Ala97, Gly98, Gln99, Gly142, Thr143. For each contiguous chain of amino acids, the peptide bond was included in the QM region and truncation occurred on the neighboring amino acid, e.g. in the case of Asp67, the carbonyl group of Val66 was included. α-tubulin amino acids included were Arg2, Asp251, Glu254, and Lys352. Each α-tubulin residue was not in a contiguous chain and was represented only by the sidechain through severing after the last CH_2_ group in the sidechain, e.g. the bond between C_β_ and C_γ_ was cut in Glu254. Every cut bond was capped with a hydrogen atom. In the case of the free heterodimer, only the β-tubulin amino acids were included. Waters in the QM region were updated every 12 hours of wall time, which ranges from 0.5 to 1 ps of simulation time. Overall, not including link atoms, the inter-dimer complex QM region consisted of 140 non-water atoms with an average of 19 waters in the region at any given time for an average total of 197 QM atoms. The free heterodimer consisted of 109 non-water atoms with an average of 27 QM waters for an average total of 190 QM atoms. The full system size for inter-dimer simulations was 182,697 atoms while for the free heterodimer the full size was 194,276 atoms with 13,519 atoms being the actual dimer (protein, GTP, Mg).

### Transition-Tempered Metadynamics Simulations

To enable the sampling of the GTP hydrolysis reaction, TTMetaD was employed. TTMetaD is a convergent form of metadynamics wherein Gaussian hills of changing height are dropped, with the hill height scaled by the smallest hill height deposited on the lowest barrier separating two basins (reactant and product).^48^ Previous work testing convergent metadynamics on the hydrolysis of ATP in actin found TTMetaD to outperform well-tempered metadynamics; however recrossings from the product to reactant well were not observed (using any form of MetaD).^28^ To increase the number of transitions observed in the present study, a multiple walker scheme was employed wherein multiple walkers add on to the same PMF.^49^ For each of the three systems studied, after an initial 50 ps seed metadynamics run, 5 pairs of walkers ran serially (pair 2 starting after pair 1 and so on) for 70 ps each for a total of 750 ps of simulation per system (and ∼10 million cpu*hours used). Pairs of walkers were launched consecutively for 70 ps each until convergence (barrier energy error < 0.5 kcal/mol, *vide infra*) was reached. Metadynamics parameters and a sample TTMetaD Plumed input section can be found in the Supplementary Information.

Two collective variables (CVs) were used to characterize GTP hydrolysis, similar to those used in previous studies of ATP hydrolysis in actin.^27,28^ The dissociation of phosphate from GDP was captured via the coordination number between P_γ_ and O_β_ (Figure 1E labels atoms involved in CVs), this CV is shown as the x-axis in Figure 2 and varies from roughly 1 in the reactant well to 0 in the product well. Coordination numbers were used instead of distances to allow ambiguity, such that P_γ_ could equally recombine with any O_β_. The association of water to P_γ_ was captured by the coordination number of P_γ_ with O_w_ and P_γ_ with O_γ_, this CV is shown as the y-axis in Figure 2 and varies from roughly 3 in the reactant well to over 4 in the product well. Additional information on the modeling of coordination numbers can be found in the Supplementary Information.

All enhanced free energy sampling was carried out using the open-source, community-developed PLUMED library (interfaced with CP2K), version 2.5.3.^50,51^ Free energy surfaces were obtained and visualized (Figure 2) with the Metadynminer R package.^52^ Minimum free energy paths (MFEPs) were obtained through the Metadynminer package with the nudged elastic band (NEB) method linking intermediate wells.^53^ Concerted pathways were obtained by NEB directly from the reactants to the products, while the dissociative pathways were formed through concatenation of two NEBs, one linking reactants to a local minimum outside of the two basins and one linking the local minimum to the products. The number of points included in each NEB for the dissociative pathway is proportional to the CV distance of the line segment connecting the two basins, with a total of 160 points for every reported pathway. In all 3 systems, one walker engaged in a phosphorylation interaction wherein the dissociating PO_3_^−^ interacts strongly with a glutamic acid residue for the remainder of the run. This was an artifact of sampling that could not lead to the product well without backtracking on the potential energy surface and was removed. This artifact and its removal is discussed in the Supplementary Information.

In order to estimate errors in the MFEPs, the collective variables for every replica were concatenated and reweighted using the final PMF, then block analysis was done. Due to the irregular shape of the sampled region of the PMF in CV space, a full averaged error would be inappropriate and instead the average error on each individual point along the MFEP was obtained from blocks containing 40,000 to 50,000 timesteps. The goal for convergence for this work was to obtain an error on the barrier of less than 0.5 kcal/mol, which was obtained for all systems after 10 walkers had reached 70 ps each.

## Data Availability

All simulation trajectories are available upon reasonable request to the authors.

## Acknowledgments

This research was supported by the National Institute of General Medical Sciences (NIGMS) of the NIH through Fellowship F32 GM140646 (D.B.) and by the U.S. Department of Energy (DOE), Office of Science, Basic Energy Sciences (BES), under Award DE-SC0023318 (G.A.V) (for latter phase methods and analysis). Computational resources were provided by the University of Chicago Research Computing Center and the U.S. Department of Defense High Performance Computing Modernization Program.

