## Supporting Information for "Unveiling the Catalytic Mechanism of GTP Hydrolysis in Microtubules"

### Supplementary Information for
Unveiling the Catalytic Mechanism of GTP Hydrolysis in Microtubules

Daniel Beckett and Gregory A. Voth
Department of Chemistry, Chicago Center for Theoretical Chemistry, Institute for Biophysical Dynamics, and James Franck Institute, The University of Chicago, Chicago, IL 60637
*Gregory A. Voth

**This PDF file includes:**

Supplementary Methods
Supplementary Analysis of Results
Sample Plumed Input File
Figures S1 to S11
Legends for Movies S1 to S8

**Other supporting materials for this manuscript include the following:**

Movies S1 to S8

#### Supplementary Methods

**Metadynamics Parameters.** Initial hill height, $w_0$, was 1 kcal/mol with a width of 0.02 coordination number units. The pace of hill deposition was every 40 timesteps or 20 fs. The bias factor, $\Delta T$, was 1240 K with a threshold bias of 1 kcal/mol to delay tempering until 1 kcal/mol worth of bias was deposited in the transition region. A tempering delay was used due to the possibility of multiple pathways and to delay any hill height decrease if a less energetically favorable crossing happened to occur first. To use TTMetaD, two basins must be defined. The reactant basin was defined as centered at (1.0, 3.2) and the product basin was defined as (0.1, 4.3). These basin centers only determine when a trajectory has crossed into the products or reactant well and are the endpoints for the paths connecting the two wells, while the only input into tempering in TTMetaD is the minimum bias on the maximally biased path between the two wells, $V^*(s(\lambda), t)$. Tempered hill height at every step is:

$$w(t) = w_0 \exp\left[-\frac{V^*(s(\lambda), t)}{k_B \Delta T}\right]$$

With tempering delayed ($w(t) = w_0$) until $V^*(s(\lambda), t)$ > 1 kcal/mol.

**Coordination Numbers.** Coordination numbers were defined as they were in the previous studies of ATP hydrolysis in actin:

$$CN = \sum_{i \in B} \frac{1 - \left(\frac{|r_i - r_A|}{\sigma}\right)^{NN}}{1 - \left(\frac{|r_i - r_A|}{\sigma}\right)^{ND}}$$

Where the coordination number of atoms belonging to group *B* to atom *A* is computed. $\sigma = 4.5$ Bohr (2.38 Å) was used as the normalizing length, with $NN = 6$ and $ND = 12$. On the *x*-axis of the 2D free energy surfaces, atom *A* is $P_\gamma$ and the atoms in *B* are the $O_\beta$ atoms, while on the *y*-axis the atoms in *B* are the $O_\gamma$ atoms and the water molecule oxygen atoms, $O_w$.

**Removal of Phosphorylation Artifact.** As mentioned in the main text Methods, a strong association of $PO_3^-$ and β:E69 creates an artifact well that must be removed from the 2D free energy surfaces. This well is at $x$ = [0.0, 0.2], $y$ = [3.0, 3.5] and can look like a fully dissociated $PO_3^-$ intermediate on the uncorrected surface but the bonding interaction between the $PO_3^-$ and the β:E69 carboxylate group forms a metastable well that does not lead to the $H_2PO_4$ product. Figure S5 shows snapshots from the final frames of the 3 walkers that inhabit this well for the longest time. These are walker W2 in the compacted interdimer complex (which can be seen in Figure S2 and in Movie S8), walker W7 in the expanded inter-dimer complex (which can be seen in Figure S3), and walker W3 in the free heterodimer (which can be seen in Figure S4). In Figure S5, bonds are drawn for associations less than 1.8 Å in length, however in practice the distance between P and the closest O fluctuates around 1.72 Å. To quantify this phosphorylation interaction over the length of the trajectory, Figure S6 shows the coordination number of $P_\gamma$ to the β:E69 carboxylate group over the lifetime of every trajectory that entered the well, not just those that inhabited it at length. From this analysis it is clear that only the walkers displayed in Figure 5 enter into this association (with coordination numbers fluctuating around 1) while others that pass through the well are able any appreciable contact with the residue.

The depth of this "phosphorylation" well displayed in the free energy surfaces can be estimated as 14.7 ± 0.4 (compacted) or 15.7 ± 0.4 (free) kcal/mol higher than the reactant well. This estimate, however, may be subject to hysteresis as this well is avoided in other walkers after the long-time occupation of the phosphorylated state, implying an overfilling of this region of the PMF. Considering this, the phosphorylation of β:E69 appears to be highly exergonic, at least 14.7 to 15.7 kcal/mol higher than the reactant, and would not be entered into under standard reaction conditions. As shown in the main text, however, dissociative pathways for GTP hydrolysis in the free heterodimer and expanded inter-dimer complex were observed and should be quantified. To do this, and to produce the MFEPs displayed in the main text, the overfilling of the phosphorylation well must be removed. For this reason, hills deposited in the walkers shown in Figure S5 were discarded after the $P_\gamma$ to β:E69 carboxylate coordination number rose above 0.3. The 0.3 cutoff was used as it was the lowest coordination number that only the 3 walkers in Figure S5 displayed, in some cases in the expanded inter-dimer complex the product well could display a coordination number of up to 0.22 without entering into the long-time phosphorylation interaction.

#### Sample Plumed Input File

Sample input file for plumed with annotations:

```
RESTART
UNITS LENGTH=A TIME=fs ENERGY=kcal/mol
INCLUDE FILE=new-ab.txt

#######These are the definitions for the constraints
b1: COM ATOMS=CA1
b2: COM ATOMS=CA2

# inter-dimer CoM distance
abdist: DISTANCE ATOMS=b1,b2

# this is the compacted configuration
restraint: RESTRAINT ARG=abdist AT=40.507 KAPPA=59.75

######CV definitions for MetaD
P: GROUP ATOMS=13513 # Gamma Phosphorus
Ow: GROUP ATOMS=_OW_,13514,13515,13516 # QM Water Oxygens AND Gamma Oxygens - last
three numbers, replace _OW_ with waters within cutoff distance at each restart via external script

Ob: GROUP ATOMS=13510,13511,13512 # Beta Oxygens

# Coordination numbers: sigma=4.5 bohr, NN = 6, ND = 12
cb: COORDINATION GROUPA=P GROUPB=Ob R_0=2.38
cw: COORDINATION GROUPA=P GROUPB=Ow R_0=2.38
```

```
# CVs vary more smoothly with extended Lagrangian approach
ex: EXTENDED_LAGRANGIAN ARG=cb,cw KAPPA=1300,1300 TAU=31.4,31.4

######Limits on the CNs, limit to two water oxygens coordinating the P
LOWER_WALLS ARG=ex.cw_fict AT=2.5 KAPPA=5.0
UPPER_WALLS ARG=ex.cw_fict AT=5.0 KAPPA=5.0
##########

m: METAD ...
  ARG=ex.cb_fict,ex.cw_fict
# 1 kcal/mol height
  HEIGHT=1
# SIGMA is in CV (CN) space
  SIGMA=0.020,0.020
# hill addition frequency (every 20 fs)
  PACE=40
# max and min values for grid
  GRID_MIN=-1,1
  GRID_MAX=2,6
# number of bins (100 per CN in range)
  GRID_BIN=300,500
  GRID_WSTRIDE=2
# can speed up restarts but also requires grid storage/writing
#   GRID_RFILE=grid-restart.dat
  GRID_WFILE=grid-out.dat
  TEMP=310
# bias factor converts to delta T for tempering
  TTBIASFACTOR=5
# delay tempering until 1 kcal/mol of bias along path
  TTBIASTHRESHOLD=1
# 3.2 and 4.3 for cw, taken from previous work on ATP
  TRANSITIONWELL0=1.0,3.2
  TRANSITIONWELL1=0.1,4.3
  GRID_NOSPLINE
# multiple walkers line, this is the 9th walker: walker ID 8
  WALKERS_DIR=../HILLS_DIR WALKERS_RSTRIDE=40 WALKERS_N=10 WALKERS_ID=9
... m:

#write every 25 fs
PRINT ARG=abdist,ex.cb_fict,ex.cw_fict,restraint.bias,m.bias FILE=colvars.dat STRIDE=50 FMT=%8.4f
```

#### Supplementary Analysis of Results

**Analysis of Alternate Pathways.** Figures S7 and S8 display the MFEPs described in the main text as well as additional possible pathways along the free energy surfaces. In the compacted inter-dimer complex, though a dissociative mechanism was not observed in any trajectory as shown in Figure S2 and the main text, for the sake of comparison we include the nudged elastic band MFEP from using the local minimum at 0.2, 3.4 as a mid-point, this being the lowest possible pathway not navigating the concerted seam shown in walkers W0, W1, W3, W5, and W6 (Figure S2). The barrier for this pathway is 26.0 ± 0.4 kcal/mol, 2.0 ± 0.6 kcal/mol higher than the concerted route, making it unlikely and further cementing the preference of the compacted lattice for the concerted pathway.

The free heterodimer displays two minima that can be used for dissociative pathways, labeled "far" and "mid" depending on the positioning along the *x*-axis in Figure S7. The "mid" pathway corresponds to the reaction shown in main Figure 3C and walker W0, however the "far" pathway can be seen in Figure S9B and corresponds to walker W1. In this pathway, the β phosphate abstracts the lytic water proton indirectly through a second water, similar to the mechanism seen in the expanded inter-dimer complex walker W0. (Figure S4) This "far" pathway, however, is 1.8 ± 0.8 kcal/mol higher than the mid dissociative, making it within error of the similar dissociative pathway in the expanded inter-dimer complex (0.7 ± 0.8 kcal/mol). This result lends further confidence to the initially surprising result of the expanded inter-dimer complex having a higher energy hydrolysis pathway than the free heterodimer, with the water wire "mid" mechanism for the free heterodimer inaccessible to the expanded complex.

Also in Figure S9 is a second dissociative mechanism seen in the expanded inter-dimer complex (Figure S9A) corresponding to walker W1 (Figure S3). This mechanism follows the same MFEP on the 2D free energy surface but leads to a different position (lower on the *y*-axis) in the product well. In this alternate mechanism, the β-phosphate directly coordinates the lytic water rather than through a second water (panel 1). Additionally, the β:E69 carboxylate also coordinates the water though the β-phosphate abstracts the proton and deposits it on an $O_\gamma$ atom. Product does not fully cross into the lowest energy portion of the product well, as can be seen in Figure S3 W1, because the lytic water O does not associate as strongly to form $H_2PO_4$, thanks to the strong charge assisted hydrogen bonding interactions on either side of the phosphate.

In Figures S7 and S8B, concerted pathways are shown for the free heterodimer and the expanded inter-dimer complex. In both cases, these pathways are slightly higher in energy than their dissociative counterparts and right on the edge of being within error, 0.5 ± 0.5 and 0.7 ± 0.7 kcal/mol higher for the free heterodimer and the expanded inter-dimer complex respectively. Figure S10 shows the concerted mechanism for the expanded inter-dimer complex (Figure S10A) and the free heterodimer (Figure S10B). In both cases, β:E69 participates directly in hydrolysis, however there are slight differences in the mechanisms. The difficult positioning of β:E69 is highlighted when considering the return circuit of the proton to form $H_2PO_4$. The proton associated with α:E254 in the compacted inter-dimer complex (Figure 3A, panel 2, main text) is able to directly return to the rotated phosphate and associate with an $O_\gamma$. In the free heterodimer, (Figure S10B, panel 2) a water wire is required to return the proton to form $H_2PO_4^{-}$

#### Supplementary Figures

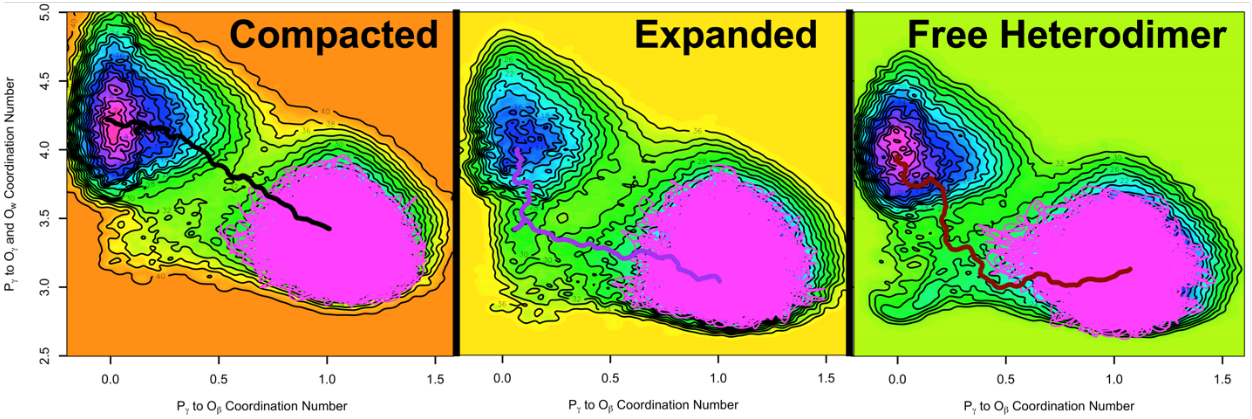


**Figure S1. CV diagrams of initial 50 ps of sampling.** Final converged free energy surfaces for GTP hydrolysis in each system (as shown in Figure 2 of the main text): compacted inter-dimer complex, expanded inter-dimer complex, free heterodimer. Lowest energy pathways shown as lines color-coded as in Figure 2 of the main text. The first 50 ps seed trajectory for each simulation is shown as a thin magenta line, located predominately in the reactant well with no transitions occurring. Probability distributions from the seed trajectories are used in Figure 4 of the main text.

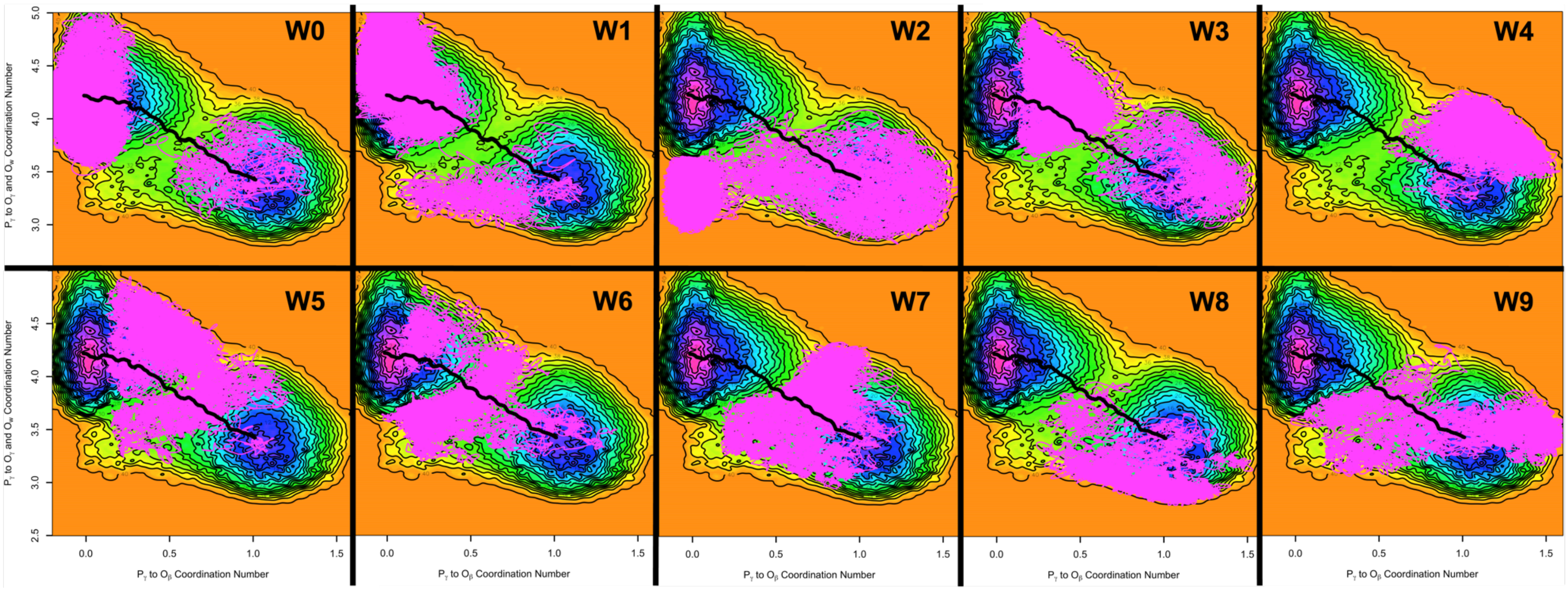


**Figure S2. CV diagrams of compacted inter-dimer complex trajectories.** Trajectory of each 70 ps walker in the compacted inter-dimer complex, walkers 0 to 9, shown as a magenta line on the final converged free energy surfaces for GTP hydrolysis. Lowest energy pathway shown as a continuous black lines. Later walkers spend the majority of time in the transition seam between the reactant and product wells and in extreme ends of either well.

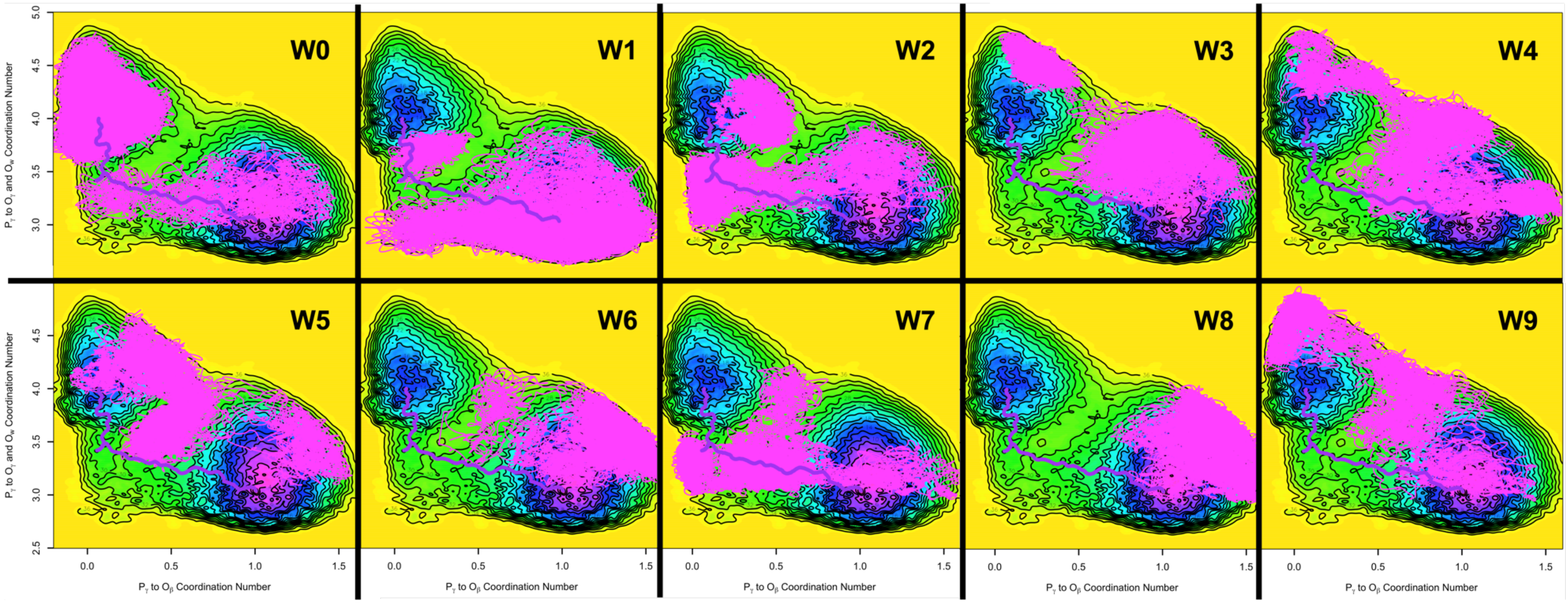


**Figure S3. CV diagrams of expanded inter-dimer complex trajectories.** Trajectory of each 70 ps walker in the expanded inter-dimer complex, walkers 0 to 9, shown as a magenta line on the final converged free energy surfaces for GTP hydrolysis. Lowest energy pathway shown as continuous purple lines. Later walkers spend the majority of time in the transition seam between the reactant and product wells and in extreme ends of either well.

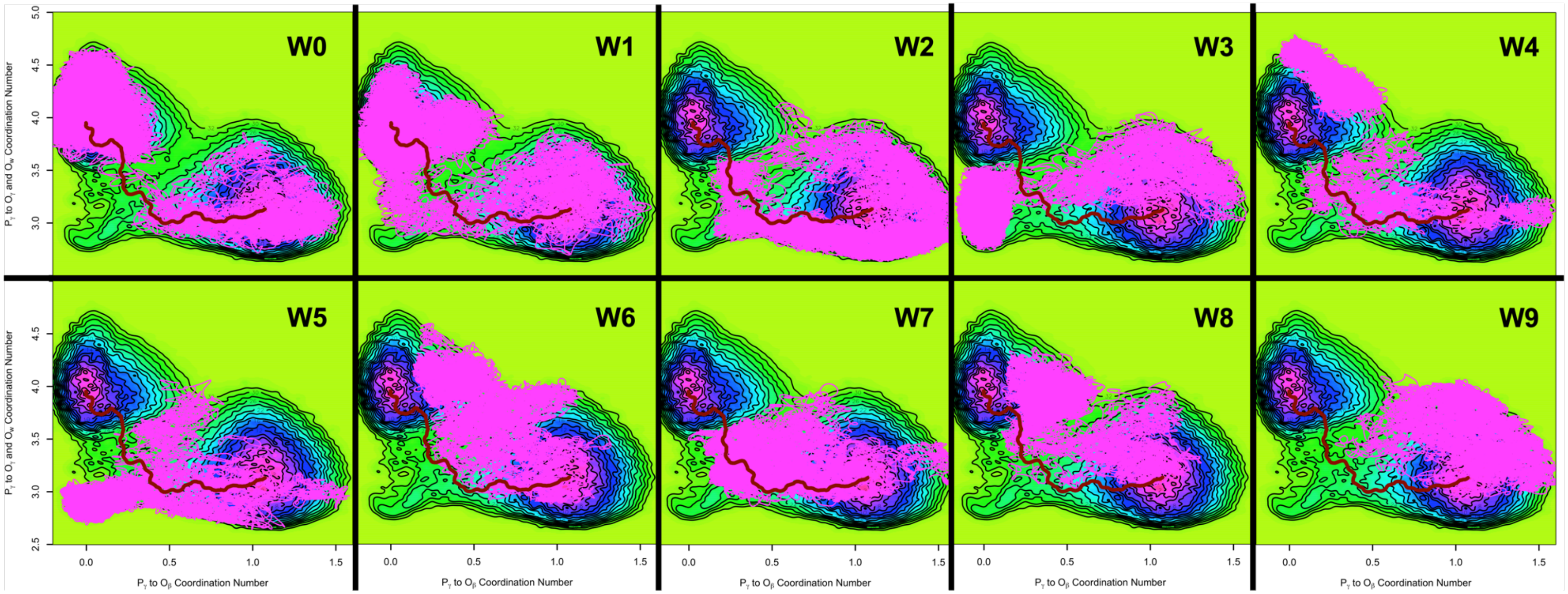


**Figure S4. CV diagrams of free heterodimer trajectories.** Trajectory of each 70 ps walker in the free heterodimer, walkers 0 to 9, shown as a magenta line on the final converged free energy surfaces for GTP hydrolysis. Lowest energy pathways shown as continuous red line. Later walkers spend the majority of time in the transition seam between the reactant and product wells and in extreme ends of either well.

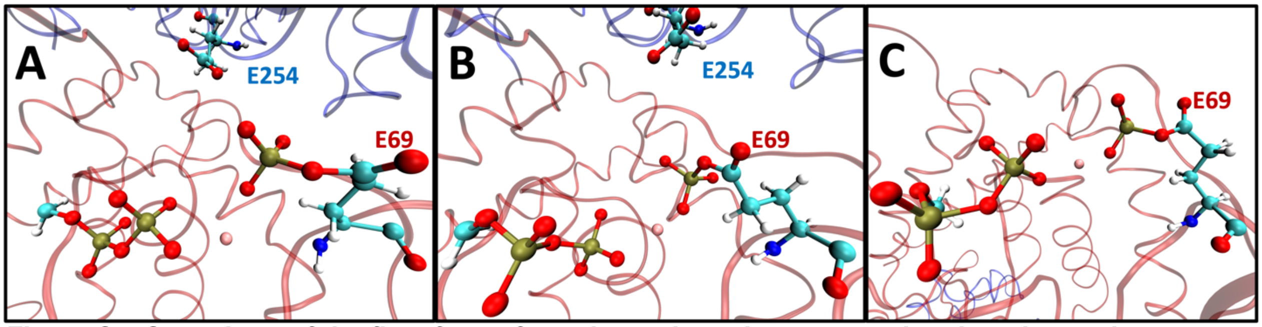


**Figure S5. Snapshots of the final frame from the walkers that occupy the phosphorylation well *x* = [0.0, 0.2], y = [3.0, 3.5] to the end of the trajectory.** A.) compacted inter-dimer complex, walker W2, B.) expanded inter-dimer complex, walker W7, C.) free heterodimer, walker W3. Trajectories of each walker can be seen in Figures S2–S4. Bonds drawn at 1.8 Å.

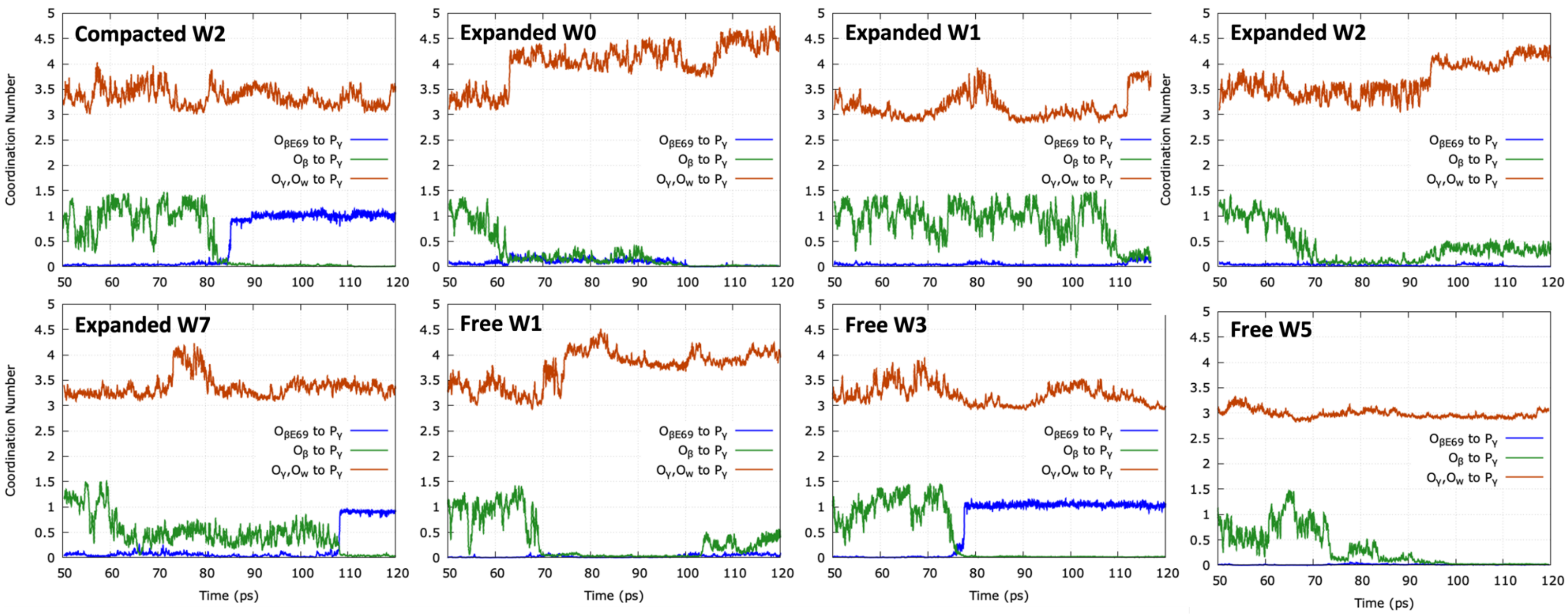


**Figure S6. Coordination number plots showing the phosphorylation artifact illustrated in Figure S5.** Coordination numbers of $P_\gamma$ to the β:E69 carboxylate group (blue) vs the reaction coordinates shown in Figure 2 of the main text ($x$ in green, $y$ in orange). Plots over the full trajectory for every walker in each system that inhabited the well $x$ = [3.0, 3.5], $y$ = [0.0, 0.2]. One replica of the compacted inter-dimer complex, four replicas of the expanded inter-dimer complex, and three of the free heterodimer. Compacted walker W2, expanded walker W7, and free heterodimer walker W3 are the only replicas with significant interaction between $P_\gamma$ and β:E69.

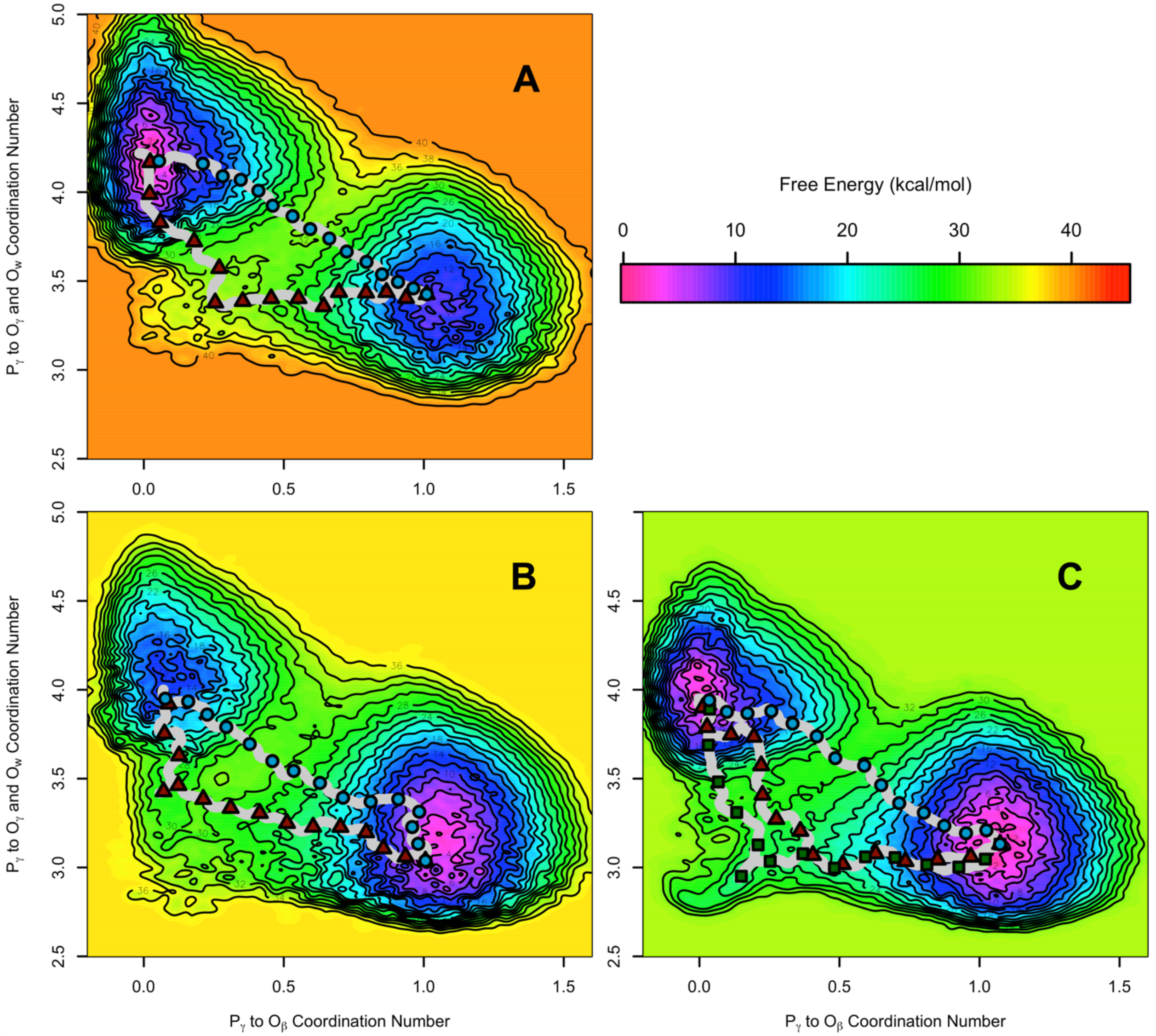


**Figure S7. All discovered nudged elastic band pathways, concerted and dissociative for each free energy surface.** A.) compacted inter-dimer complex, B.) expanded inter-dimer complex, C.) free heterodimer. Every possible low energy navigation of the free energy surface is shown: the concerted routes as described in the main text are shown with blue circles. Dissociative routes are shown with red triangles in all three systems, including compacted where a dissociative mechanism was not observed directly in a walker but can be calculated from the free energy surface. Both possible dissociative routes for the free heterodimer are shown, with the higher energy “far” route illustrated with green squares. Topological lines drawn every 2 kcal/mol. Each pathway is represented by 160 points in total, every 10$^{th}$ point represented as a visible point on the plot for comparison.

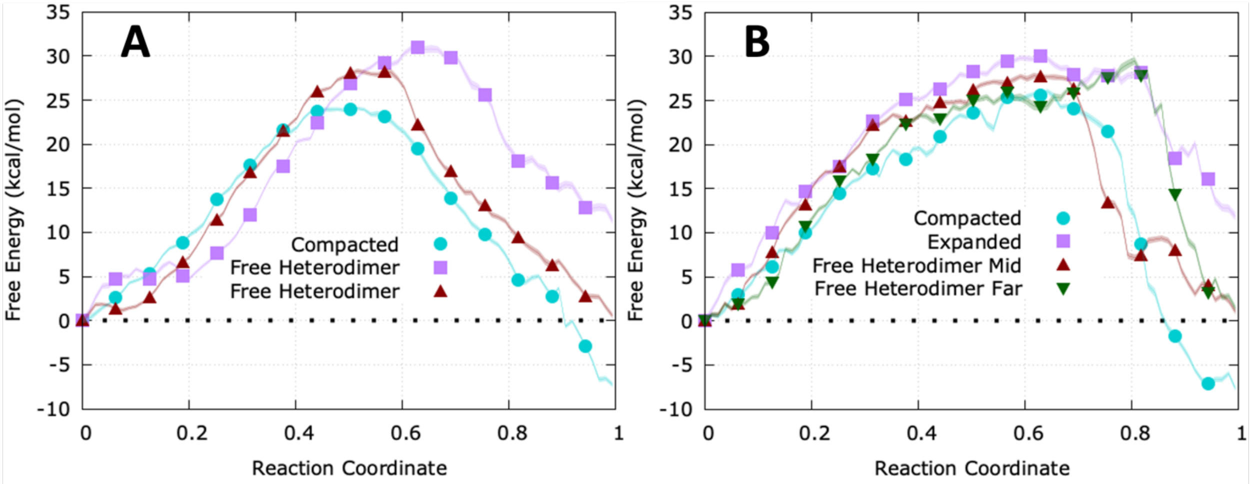


**Figure S8. Minimum free energy pathways for each system as shown in Figure S7.** A.) concerted reactions (blue circles in Figure S7) and B.) dissociative reactions (red triangles and green squares in Figure S7). Error shown as translucent region around lines, each line composed of 160 points with symbols at every 10th point.

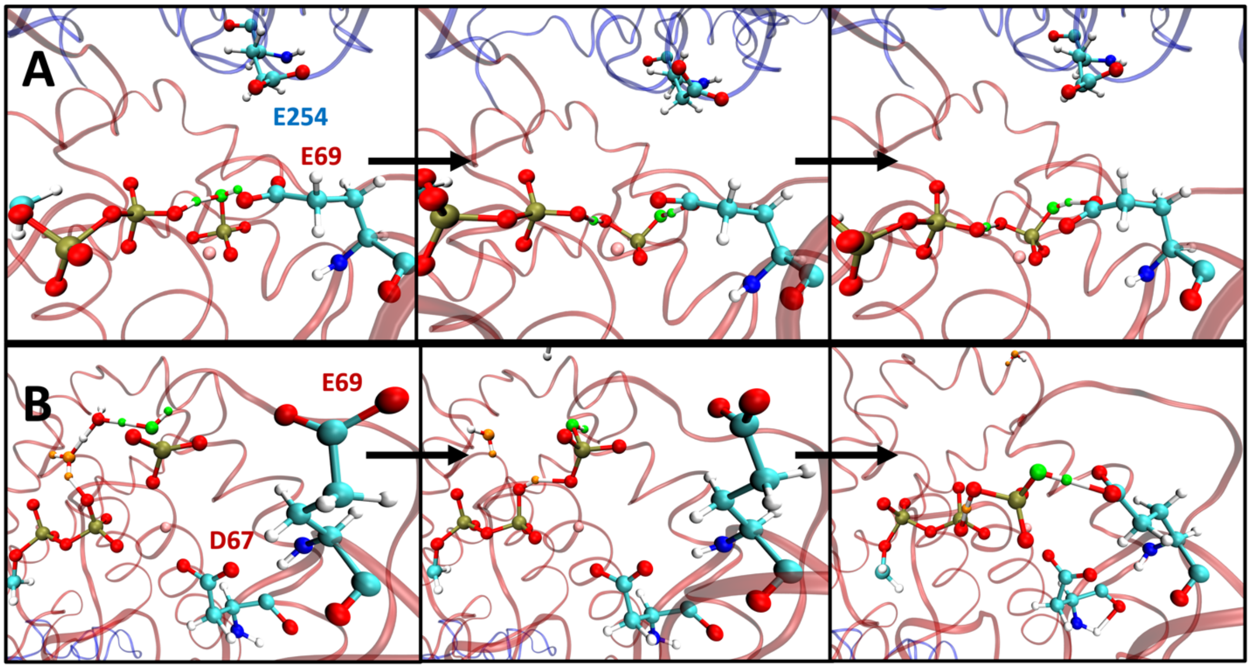


**Figure S9. Snapshots of proton rearrangements from metadynamics trajectories for alternate dissociative pathways.** Snapshots of proton rearrangements from metadynamics trajectories for dissociative reactions in A.) expanded inter-dimer complex (taken from walker W1) and B.) free heterodimer (W1). Lytic water associating with leaving phosphate is colored green, additional water depositing proton onto the phosphate is colored orange when present, magnesium shown in pink. Bonds drawn with a 1.8 Å cutoff, only critical residues shown atomically (labeled on the first panel), red ribbons represent β-tubulin while blue represent α-tubulin. The first column shows proton transfer to the catalytic residue, the second column is proton transfer from the catalytic residue to form $H_2PO_4$, and the third column shows final product after formation (or final proton transfer, B). Movies pulled directly from the appropriate trajectory for each mechanism shown here can be found in the Supplementary Information.

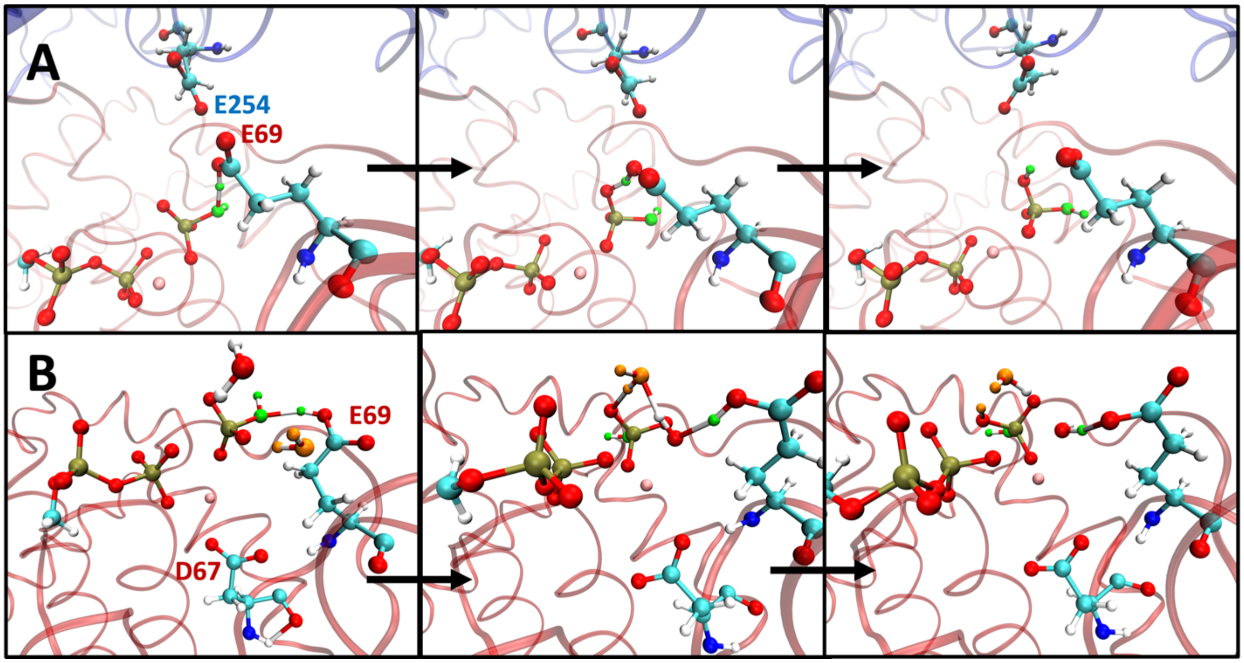


**Figure S10. Snapshots of proton rearrangements from metadynamics trajectories for higher energy concerted reactions.** A.) expanded inter-dimer complex (taken from walker W5) and B.) free heterodimer (W4). Lytic water associating with leaving phosphate is colored green, additional water depositing proton onto the phosphate is colored orange when present, magnesium shown in pink. Bonds drawn with a 1.8 Å cutoff, only critical residues shown atomically (labeled on the first panel), red ribbons represent β-tubulin while blue represent α-tubulin. The first column shows proton transfer to the catalytic residue, the second column is proton transfer from the catalytic residue to form $H_2PO_4$, and the third column shows final product after formation. Movies pulled directly from the appropriate trajectory for each mechanism shown here can be found in the Supplementary Information.

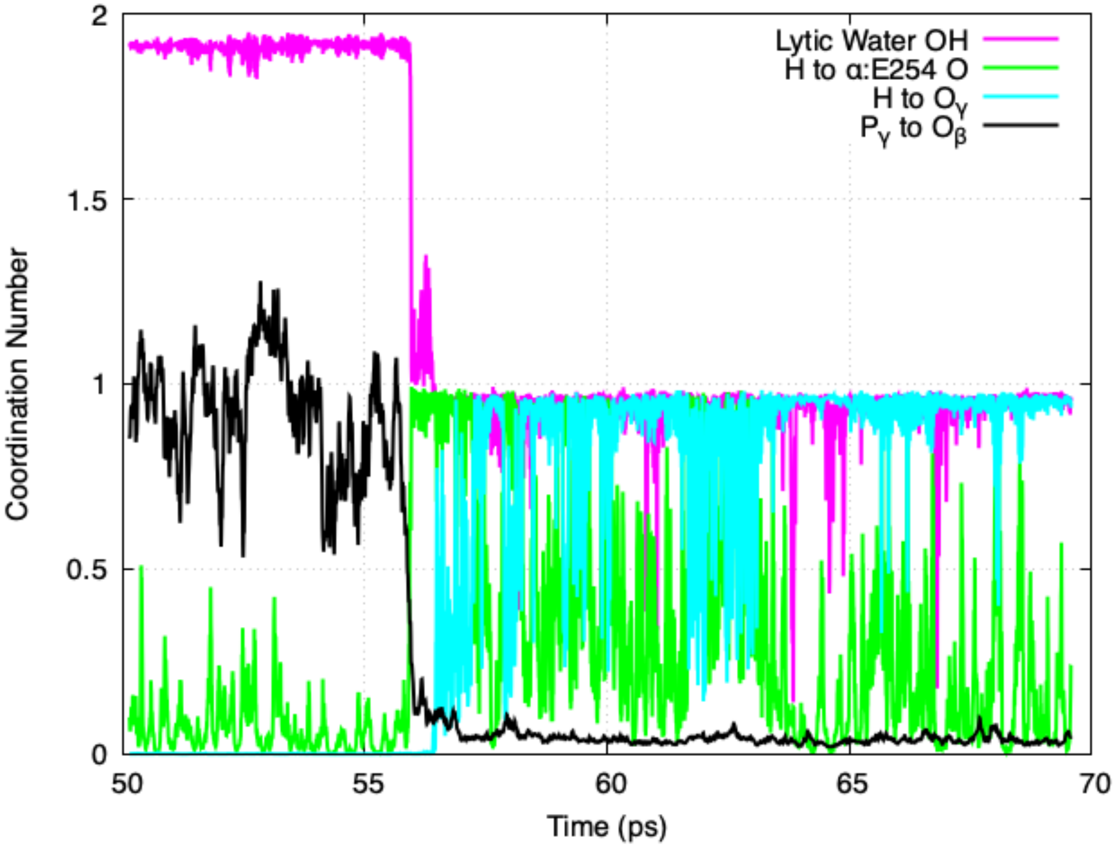


**Figure S11. Example of critical coordination numbers for hydrolysis in the compacted inter-dimer complex (walker W0).** The first 20 ps of walker W0 in the compacted inter-dimer complex with respect to coordination numbers of the lytic water hydrogens to the lytic water oxygen (magenta), lytic water hydrogen to the α:E254 residue oxygens (green), lytic water hydrogen to γ-phosphate oxygens, and the *x*-axis coordinate of the 2D PMFs (coordination number of γ-phosphate phosphorus to β-phosphate oxygens. Dissociation of the γ-phosphate is coupled to proton abstraction by α:E254 and the proton is quickly shuttled to a γ-phosphate oxygen, after which the $H_2PO_4$ product interacts with α:E254 for ~10 ps.

#### Legends for Movies

Each movie is a snapshot of the hydrolysis reaction pulled from the indicated trajectory at the indicated timesteps. Bonds are drawn when contacts are less than 1.8 Å apart. The water adding to the releasing phosphate is colored in green while, if present, the water donating the final proton for the formation of $H_2PO_4$ is colored orange. Each frame of video is 50 fs of simulation.

**Movie S1:** GTP hydrolysis in the compacted inter-dimer complex following the concerted route, 50 ps to 80 ps of Walker W0. Snapshots shown in main text Figure 3A.

**Movie S2:** GTP hydrolysis in the expanded inter-dimer complex following the concerted route, 50 ps to 90 ps of Walker W5. Snapshots shown in Figure S10A.

**Movie S3:** GTP hydrolysis in the free heterodimer following the concerted route, 60 ps to 90 ps of Walker W4. Snapshots shown in main text Figure S10B.

**Movie S4:** GTP hydrolysis in the expanded inter-dimer complex following a dissociative route, 60 ps to 105 ps of Walker W0. Snapshots shown in main text Figure 3B.

**Movie S5:** GTP hydrolysis in the expanded inter-dimer complex following a dissociative route, 90 ps to 120 ps of Walker 1. Snapshots shown in Figure S9A.

**Movie S6:** GTP hydrolysis in the free heterodimer following the lowest energy dissociative route, 50 ps to 85 ps of Walker 0. Snapshots shown in Figure 3C.

**Movie S7:** GTP hydrolysis in the free heterodimer following the higher energy dissociative route, 60 ps to 80 ps of Walker 1. Snapshots shown in Figure S9B. Illustrates initial proton transfers, final return circuit to form $H_2PO_4$ (Figure S9B, Panel 3) begins late in the trajectory (~115 ps).

**Movie S8:** Phosphate dissociation in the compacted inter-dimer complex, 80 to 120 ps of Walker 2 and the formation of the artifact well. Snapshot shown in Figure S5A.
